# Parent-offspring conflict over sex determination in non-Mendelian systems

**DOI:** 10.1101/2025.11.26.690632

**Authors:** Thomas J. Hitchcock, Robert B. Baird

## Abstract

Across the tree of life, many organisms exhibit asymmetric inheritance systems in which males and females contribute differently to the long-term genetic future of the population. Although, in such groups, the sexes ultimately differ in their contributions, the zygotes that become males and females often start out genetically identical, with sex determined by maternal factors deposited into the embryo. However, there has been little work considering what the optimal sex ratio is from the perspective of the offspring in such scenarios, how this may differ from their parents, and how such conflicts may be modulated by other ecological factors. To investigate this, we develop analytical models to calculate the optimal sex allocation under a range of asymmetric genetic systems, and under the control of different parties. We then investigate the effects of various population structures and mating systems to consider their effects in shaping such conflicts. We find that asymmetric genetic systems may be prone to perpetual ongoing conflict between mothers and offspring over sex determination, even in panmictic populations. This may be one factor explaining the diverse and unusual sets of sex determining systems seen in these groups.

## Introduction

Conflicts over the sex ratio are among the most well-characterised examples of evolutionary conflict (West 2009). Different elements - individuals, chromosomes, symbionts, or individual loci - can all exhibit different optima for the sex ratio, as they may be transmitted differently through males and females, and/or differ in their relatedness to social partners (Burt and Trivers 2006). The coevolutionary struggles between these different parties can result in overhauls in sex determination mechanisms, and is thought to be one of the reasons that sex determination systems are so diverse (Bull 1983; Beukeboom and Perrin 2014).

The genetic system of the focal organism shapes sex ratio conflict in two fundamental ways. First, by defining how genes pass from mothers and fathers to sons and daughters, it determines the relative value of producing a male versus a female for each party. Second, it constrains when and how individuals can exert control over sex allocation. For example, in the most common asymmetric genetic system, arrhenotoky, females are produced from fertilised eggs and males from unfertilised ones. As males and females are born with different ploidies, it is then often assumed that offspring cannot directly influence their own sexual fate. For a similar reason, it has typically been assumed that fathers have little influence over sex allocation in arrhenotokous species (although see Shuker et al. 2006; King et al. 2012; Macke et al. 2014).

However, there are many organisms where genetic asymmetries emerge after, not at, fertilisation. For example, in species with paternal genome elimination (PGE) (Normark 2003; Ross et al. 2022; Figure 1), both males and females are initially born with the same symmetric complement of genes from mothers and fathers. Only later, after sex determination has occurred, do asymmetries emerge; with males eliminating their paternal-origin genome, and thus only transmitting their maternal-origin chromosomes. Whilst mothers may still play an outsized role – as the early stages of embryogenesis tend to be maternally controlled (Tadros and Lipshitz 2009) – both the genome of the father, and of the offspring, may be able to play important roles in modulating the sex determination cascade in such species. Consequently, asymmetric systems such as PGE reopen the possibility of paternal and offspring agency in sex allocation decisions.

**Figure 1.**
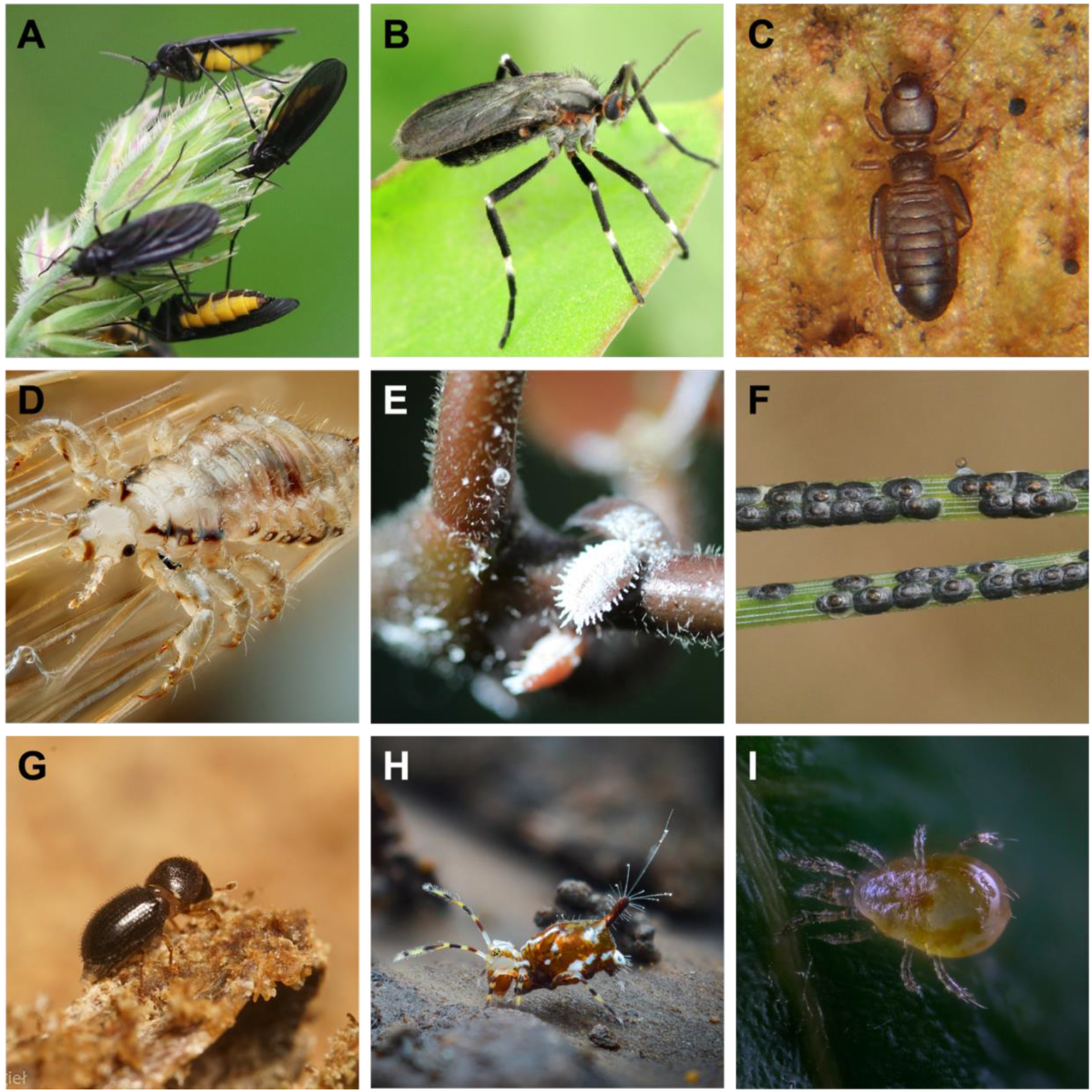
Known systems with paternal genome elimination (PGE). (A) Dark-winged fungus gnats (Sciaridae, credit: Charlie Farrell, CC0). (B) Gall midges (Cecidomyiidae, source: Stephen Luk, CC-BY-NC, iNaturalist: 102505337). (C) Booklice (Psocodea, source: Alexei Kouprianov, CC-BY-NC, iNaturalist: 267411141). (D) Parasitic lice (Phthiraptera, source: Giles San Martin, CC-BY-SA, Flickr: 4900275659). (E) Scale insects (Neococcidae, source: Scott F Smith, CC-BY-NC, iNaturalist: 259770597). (F) Armoured scale insects (Diaspididae, source: James Bailey, CC-BY-NC, iNaturalist: 109351010). (G) Coffee berry borer beetle (*Hypothenemus*, source: Hubert Szczygieł, CC-BY-NC, iNaturalist: 195150681). (H) Globular springtails (Symphypleona, source: Danny Radius, CC-BY-NC, iNaturalist: 260512611). (I) Predatory mites (Phytoseiidae, source: Dokhtoruk Andrii, CC-BY-4.0, Wikimedia Commons). Images have been cropped.

Despite this, there has been little theoretical work considering how conflict between parents and offspring may play out in such asymmetric systems. Two notable exceptions are the works of Bull (1983) and Haig (1993). Bull modelled the evolution of the sex ratio under PGE when under zygotic control, finding that the sex ratio evolved to a value of ⅓, although he stated that the intuition behind this result was unclear (1983, Chapter 13). Haig (1993) also considered the optimal sex ratio from the perspective of the maternal-origin and paternal-origin genomes in the zygote under PGE, but assumed that these were identical to the interests of the parents they descended from, and thus used Hamilton’s previous derived expressions for X chromosomes (Hamilton 1979). In addition, other authors have considered how mating systems and demography might generate conflicts between parents and offspring over the sex ratio (Trivers 1974; Werren and Hatcher 2000; Pen 2006; Wild and West 2009). Wild and West in particular considered both imprinting and haplodiploidy, but their analysis assumed that haplodiploid workers controlled the sex of offspring, rather than the genomes of the zygote directly. How these different results connect, and thus the interplay between the asymmetric genetics, demography, and parent-offspring conflict remains unclear.

Here, we develop a model of sex allocation within an asymmetric genetic system where an individual’s sex may be under the control of its mother, its father, or the individual itself as a zygote. We allow both the patterns of parental transmission and zygotic expression to vary so that we can uncouple their contributions and explore a range of possible genetic systems. We provide an intuitive explanation of the origin of Bull’s ⅓ result, and how this changes depending on the genetic system and expression patterns in early development. We extend this to consider the effects of local mate competition and resource competition, under a range of different mating systems, and compare our results to previous models of parent-offspring conflict under population structure. Finally, we discuss how our current understanding of sex determination in these groups might hint at a history of parent-offspring conflict over the sex ratio.

## Methods

### Genetics and life cycle

We begin by describing the genetic parameters that underlie our model. We assume that all offspring are initially diploid, inheriting one gene copy from their mother and one from their father. The phenotype an offspring expresses is determined by the relative influence of these two genomes. Let *ρ* denote the proportion of control exerted by the maternal-origin genome; the paternal-origin genome then contributes the remaining 1 − *ρ*. An individual’s sex ratio phenotype can therefore be written as:

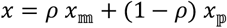

where *x*_𝕞_ and *x*_𝕡_ represent the genetic values of the maternal- and paternal-origin genomes, respectively.

Juveniles undergo sexual differentiation, developing as males with probability *x* and as females with probability 1 − *x*. After differentiation, the two genomes within an individual may be transmitted to future generations with different probabilities. A female gamete carries the maternal-origin copy with probability *α* and the paternal-origin copy with probability 1 − *α*. Conversely, a male gamete carries the paternal-origin copy with probability *β* and the maternal-origin copy with probability 1 − *β*. These genetic processes are illustrated in Figure 2.

**Figure 2.**
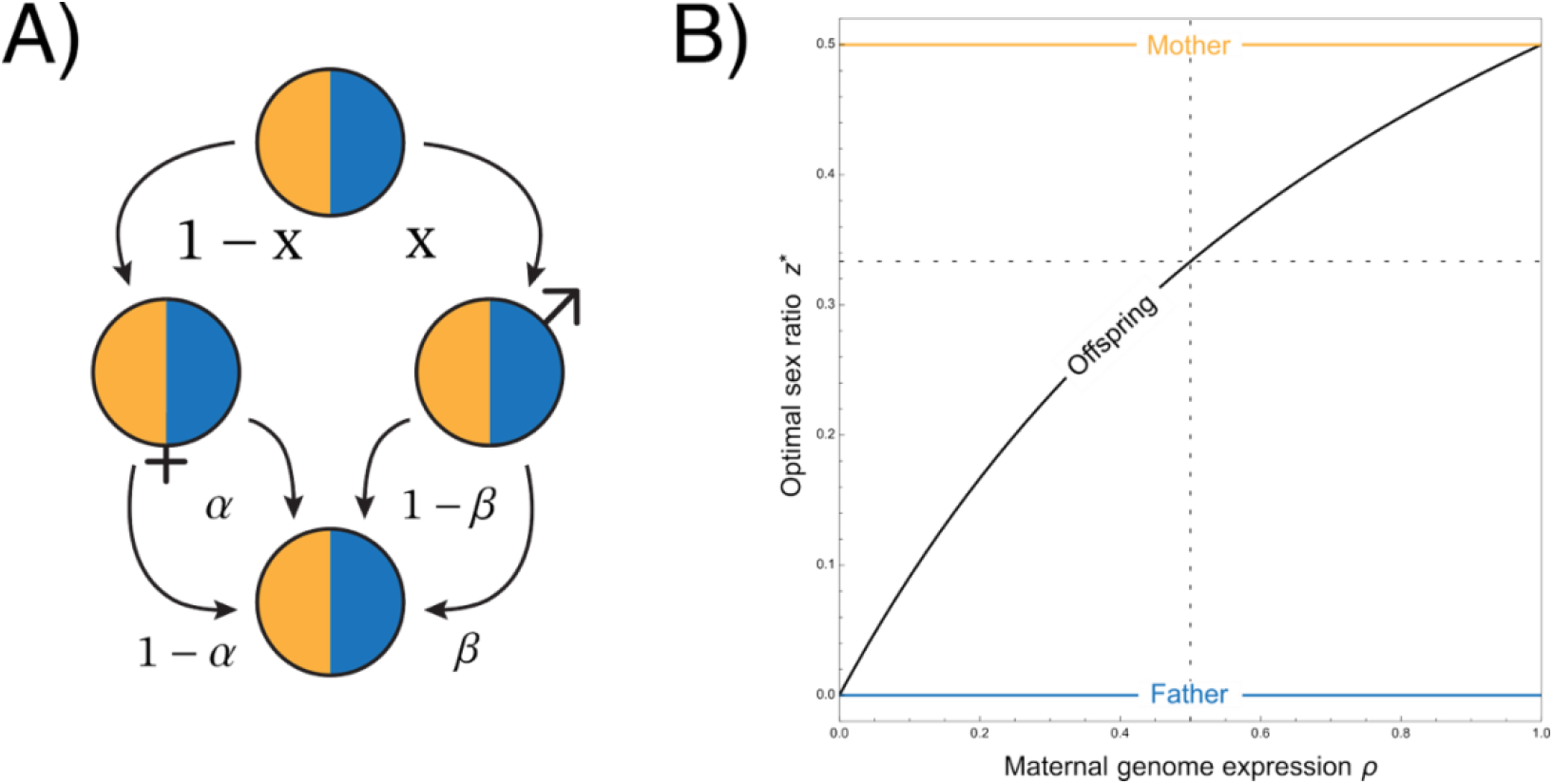
a) Schematic of inheritance system and b) relative control over zygotic sex ratio phenotype and the optimal sex ratio under PGE (*α* = 1/2, *β* = 0).

We embed this genetic system within the following life cycle. The population consists of a large (effectively infinite) number of patches, each containing *N* mated females. The life cycle proceeds as follows. (1) Birth: each female produces a large number of offspring on her patch before dying. (2) Sexual differentiation: juveniles commit to becoming either males or females according to their phenotype *x*. (3) Dispersal and mating: offspring disperse from their natal patch with sex-specific probabilities *d*_*f*_ for females and *d*_*m*_ for males. Mating occurs after male dispersal, and either before female dispersal (DDM life cycle) or after female dispersal (DMD life cycle) (Wild & Taylor 2004; Supplementary Figure 1). (4) Competition: mated females compete for the *N* breeding positions available on each patch.

### Relatedness coefficients and reproductive values

Given the genetic system and demography described above, we next compute the genetic correlations between different gene sets at different points in the life cycle – the relatedness coefficients. We denote the relatedness between two genetic states 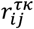: *i* and *j* refer to the parental origin of the two genes (maternal, 𝕞, or paternal, 𝕡), *τ* is the controlling actor (zygote (*Z*), mother (*M*), or father (*F*)), and *κ* refers to whether the recipient is oneself (or own offspring) *S*, or another member of the patch *P*. For example, the relatedness between the maternal-origin gene in a mother and the paternal-origin gene in the juveniles on the patch would be 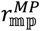. Full methods for calculating the relatedness coefficients can be seen in the Supplementary Material (SM§2).

The genetic system will also alter the expected long term contributions that different classes of genes make to future generations - i.e. their class reproductive values (Fisher 1930; Price and Smith 1972; Taylor 1990). We write the reproductive value of maternal-origin genes as *c*_𝕞_, and the reproductive value of paternal-origin genes as *c*_𝕡_. These are obtained as the stationary distribution of the gene transition matrix: *T* = (*t*_*ij*_), where *t*_*ij*_ is the probability that a gene in state *i* ∈ 𝕞, 𝕡 descends from a gene in state *j* in the previous generation. Solving yields:

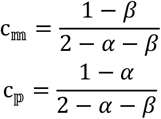

We can also directly relate this to the reproductive values of males and females as classes. As females contribute only to maternal-origin genes, and males only to paternal-origin genes then the class reproductive value of females is equivalent to the maternal-origin genes *c*_*f*_ = *c*_𝕞_ and the class reproductive value of males is equivalent to the paternal origin genes *c*_*m*_ = *c*_𝕡_.

### Inclusive fitness

Let the population be monomorphic for a sex ratio *z* (the probability of developing as a male). Consider a rare mutant that perturbs the genetic value by a small additive amount *x* = *z* + *δ*. In the case of our maternal origin copy, this will cause a phenotypic change of *ρδ*, and for our paternal-origin copy a phenotypic change of (1 − *ρ*)*δ*. We then seek an evolutionary stable sex ratio strategy, i.e. one in which a mutant strategy cannot invade the resident (Christiansen 1991; Taylor 1996). In order for this mutant strategy to invade the population it must on average increase the inclusive fitness of its bearer (Hamilton 1964; Taylor 1996; Scott & Wild, 2023). Let us write the change in the inclusive fitness of our actor *i* as Ω_*i*_. If the same locus underpins the phenotype of multiple actors, then we must sum the inclusive fitness effects over these different actors that our mutant may be expressed in 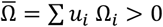, where *u*_*i*_ is the relative abundance of state *i* in the population. There are two states that our mutant may find themselves in, a maternal-origin state or a paternal-origin state. In our model, they compose equal fractions of the population. We can then sum the inclusive fitness effects experienced across these states, by the fraction of the population in this state, and its relative phenotypic effects when expressed here. Thus:

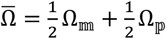

A fuller expression of the different components of this expression can be seen in the supplementary material (SM§2). We can then solve for when the inclusive fitness change will be Ω = 0, and then numerically check that this is a convergent singular strategy.

## Results

### Panmixia

First, we focus on the case of panmixia and thus assume that the two gene copies within an individual are unrelated. Our change in inclusive fitness can be written as:

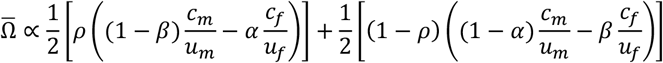

Which by solving for when 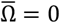, and rearranging, gives us an optimal sex ratio of:

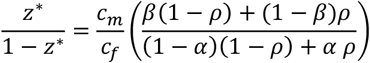

When the maternal-origin and paternal-origin genomes exert equal control over the sex ratio phenotype, *ρ* = 1/2, then a gene’s optimal investment in sons equals their population-level reproductive value, so that the probability of producing a male zygote equals the class reproductive value of males, *z*^∗^ = c_m_. In the case of PGE (*α* = 1/2, *β* = 0), then this reduces to *z*^∗^ = 1/3, the value originally obtained by Bull (1983).

However, this result only holds when those two genomes have equal control over the phenotype. When the maternal-origin genome exerts full control (*ρ* = 1) then the optimal sex ratio is the share of the maternal-origin gene’s reproductive value that comes from males *z*^∗^ = ((1 − *β*)*c*_*m*_)/c_𝕞_. Equally, when it is paternally-controlled (*ρ* = 0), then the optimal sex ratio will be the share of the paternal-origin gene’s reproductive value that comes from males *z*^∗^ = (*β c*_*m*_)/c_𝕡_. In this way, these genomes exactly reflect the interests of their parents. For the case of PGE, we can see that this means the maternal-origin gene in the zygote (and thus the mother) should favour a sex ratio of *z*^∗^ = 1/2, whilst paternal-origin copy (and father) favours a sex ratio of *z*^∗^ = 0 (Haig 1993) (Figure 2B).

### Local mate competition

Males and females may interact with relatives, impacting the inclusive fitness valuations of being a male or a female. Classically, Hamilton showed that if males compete locally, but females globally, then this may favour mothers to produce female-biased broods (Hamilton 1967). Trivers and others showed that whilst a similar force acts on an offspring’s decision, they typically favour less biased sex ratios, as they benefit from being the rarer sex, even at a cost to total local productivity (Trivers 1974; Werren & Hatcher, 2000; Pen 2006; Wild & West 2009). This can generate conflicts between parents and offspring over such sex allocation decisions.

To incorporate such scenarios, we now consider local mate competition scenario, where males remain on their natal patch *d*_*m*_ = 0, but females disperse fully after mating *d*_*f*_ = 1. This is the same scenario analysed by Hamilton (1967, Hamilton 1979) and corresponds to the dispersal-mating-dispersal (DMD) life cycle described earlier. Solving again for the equilibrium condition 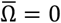, we can find the optimal sex ratio *z*^∗^. Focusing on monogamy and PGE under equal control, the optimal sex ratio from the zygote’s perspective is:

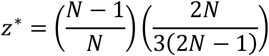

As the patch size, *N*, decreases, offspring favour an increasingly female-biased sex ratio. When *N* is large, this is more female-biased than the sex ratio favoured by mothers 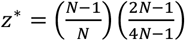, but as *N* gets smaller, the situation reverses, with offspring favouring a more male-biased sex ratio that their mothers when the patch size is small (*N* ≈ 2). This reversal reflects a blend of effects discussed above – offspring value being the rarer sex but also place more value on female fitness. As with panmixia, this also reflects an averaging over the maternal-origin and paternal-origin interests. The maternal-origin genomes favour a more male-biased sex ratio, and paternal-origin genomes favouring a more female-biased sex ratio.

Additionally, there is not only strong conflict between mothers and offspring, but between fathers and mothers too. Solving the same scenario under paternal-control, the optimal sex ratio is:

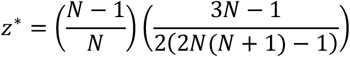

As fathers only directly transmit genes through daughters, then it is only through these individuals that they gain direct fitness, and thus favour a more female-biased sex ratio than either zygotes or mothers. However, as inbreeding increases, they become increasing related to their sons, and so place more value on the production of males. Importantly, as with zygotic control, there is also strong conflict here between the interests of the paternal-origin and maternal-origin genomes. In an outbred population the interests of the paternal-origin genome in fathers have no impact, as any change in sex allocation will not impact its inclusive fitness. But with population structure, these genomes may be related to their patchmates, and thus changes in sex allocation will alter their inclusive fitness. Whilst a male’s maternal-origin genome favours a highly female-biased sex ratio, the paternal-origin genome favours a relatively more male-biased sex ratio (Figure 3a). This means that males under PGE will favour a less extreme female bias than their arrhenotokous counterparts. Full expressions for this, and other mating schemes, can be seen in the SM§5.

**Figure 3.**
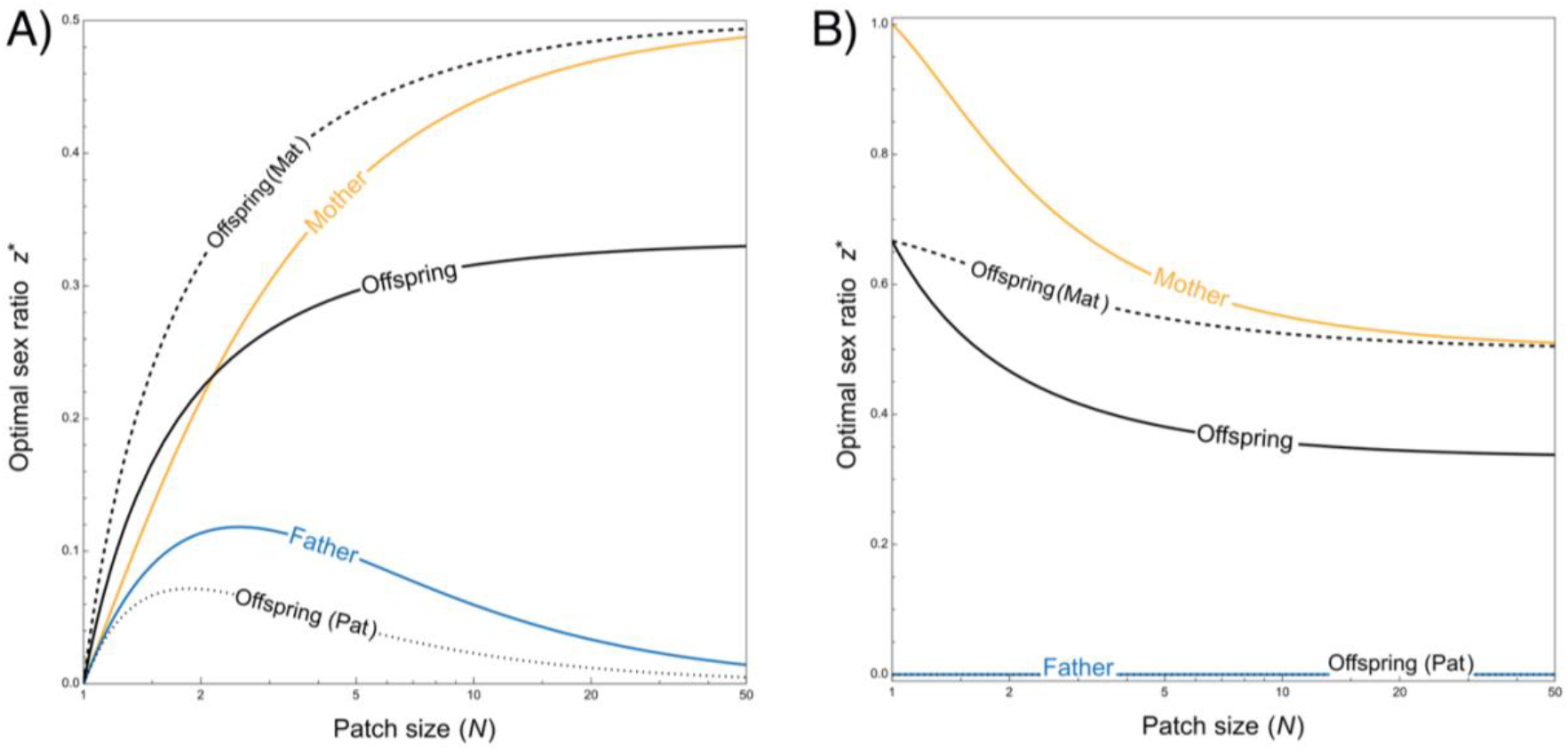
Optimal sex ratio for different actors under paternal genome elimination (PGE) and monogamy, either in a scenario with (A) local mate competition, and (B) local resource competition.

### Local resource competition

There may be scenarios where, rather than competition between brothers, the dominant ecological factor is instead competition between related females. This is commonly known as local resource competition (Clarke 1978). In such scenarios, if females instead compete locally for resources, whilst males compete globally for matings, then a male-biased sex ratio may be favoured. We analyse such a scenario, first considering monogamy, and PGE. Under offspring control, the optimal sex ratio will be:

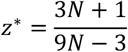

When the patch size is very large (*N* → ∞), then offspring favour a sex ratio of 1/3 as before. As patch size decreases, then they favour the production of more males, although they only favour a strictly male-biased sex ratio when the patch size is very small (*N* ≈ 1.5). Notably zygotes always favour a more female-biased sex ratio than their mothers, 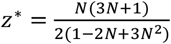 (Wild & West 2009). As before, there is also strong conflict between maternal-origin and paternal-origin genomes within the zygotes in this scenario. Whilst the maternal-origin genomes qualitatively follow the above-described pattern, the paternal-origin genomes only favour the production of daughters. Thus, unlike the local mate competition scenario, in this local resource scenario the conflict between these genomes increases, rather than decreases, as the patch size decreases (Figure 3b).

Fathers under local resource competition favour the production solely of daughters *z*^∗^ = 0. Even as the patch size decreases, because males disperse fully, the patch size does not change their relatedness either to their offspring or other members of the match. Consequently, unlike, with local mate competition, the magnitude of conflict between mothers and fathers increases as the patch size decreases. Once again, full expressions for some other mating systems, and comparisons to previous results can be seen in SM§5.

## Discussion

Parent-offspring conflict over sex allocation has traditionally been explored in the context of symmetric inheritance systems, with sex governed by the random segregation of XY or ZW chromosomes. In such scenarios, under random mating, an optimal sex ratio of ½ is generally aligned between parents and their offspring (Cobb 1914; Fisher 1930; Gardner 2023). However, in asymmetric systems such as paternal genome elimination (PGE), where males inherit but do not transmit their paternally inherited chromosomes, this alignment breaks down. Previously, Bull (1983) predicted that, in such systems, offspring should favour a biased sex ratio of ⅓ males, while mothers still favour ½. Here we demonstrate that this difference can be understood as a balance between the inclusive fitness effects of maternal-origin and paternal-origin chromosomes: the former benefit from being transmitted through both sons and daughters, while the latter are only transmitted through daughters. However, we also show that this ⅓ rule does not hold universally, as it is sensitive to biases in the expression of the maternally versus paternally inherited alleles, as well as non-random mating or population structure.

This divergence in sex ratio optima creates potential for an evolutionary arms race over control of sex determination to ensue. Since early zygotic development is governed first by maternally deposited transcripts, and then by zygotically expressed genes, with sex determination ultimately being controlled by an interaction between the two (Harrison et al. 2023), this is most likely to play out between the mother and zygote. Paternally-inherited alleles, if they are not silenced, may serve as zygotic signals that could influence sex determination following zygotic genome activation. Such zygotic control could manifest as sex-specific transcriptional regulation or the evolution of male suicide alleles, where paternally inherited alleles promote their own elimination to skew the sex ratio towards females (Ross et al. 2011) (although this will depend on patterns of reproductive compensation (Hastings 2001)). In response, selection should favour the evolution of maternal suppressors. In scale insects, coffee berry borer beetles, and booklice, which all have PGE, the paternal genome is transcriptionally silenced in early embryogenesis (Nur 1980; Brun et al. 1995; Hodson et al. 2017), which may represent an outcome of such conflict.

In almost all of the systems so far characterised (Figure 1), maternal control over sex determination appears to dominate. Mothers are able to influence the sexual karyotypes of offspring, which they do via embryonic elimination of chromosomes in the case of fungus gnats (Metz 1938), gall midges (Stuart and Hatchett 1991), armoured scale insects (Brown 1965), predatory mites (Nelson-Rees et al. 1980), and globular springtails (Dallai et al. 2001), or embryonic silencing of chromosomes in booklice (Hodson et al. 2017), scale insects (Nur 1980), and coffee berry borer beetles (Brun et al. 1995). In all these cases, the result is that the affected embryos become haploid, functionally haploid (via silencing), or haploid for the X chromosome, which results in them developing into males. If these represent cases where mothers have ‘won’ the conflict over sex determination then this is perhaps unsurprising, since maternally-deposited factors control early embryonic development. However, this does not preclude the zygote from influencing sex determination by targeting downstream sex determination genes. Across many insects, the canonical *transformer*-*doublesex* pathway initiates sexual development (Verhulst et al. 2010). While *transformer* is rapidly evolving and appears to be absent from many lineages (Geuverink and Beukeboom 2014), alternative splicing of the downstream transcription factor *doublesex* into male- and female-specific isoforms is virtually ubiquitous across insects (Shukla and Nagaraju 2010; Wexler et al. 2019). It is then highly unusual that in three of the PGE clades listed above, fungus gnats (Ruiz et al. 2015), scale insects (Bain et al. 2021), and parasitic lice (Wexler et al. 2019), *doublesex* is not alternatively spliced into male- and female-specific transcripts (Table 1). It is tempting to speculate that this pattern could represent a relic of zygotic attempts to override sex determination, but further work exploring the sex determination mechanisms across PGE systems will be required to explore this hypothesis.

**Table 1.** In the systems with paternal genome elimination (PGE), timing of PGE varies between systems, but with the common result that males fail to pass on their paternally-inherited chromosomes. In most of these systems, mothers are known to control offspring sex through chromosome elimination or silencing. Interestingly, in three of the systems, the downstream canonical sex determination transcription factor *doublesex* does not follow the orthodox pattern seen in virtually all other insects. Note that gall midges are reported to have sex-specific transcripts, but this result was in pupae and was in a non-peer reviewed thesis, so it was not included in the table below. For a more comprehensive overview of PGE systems and their mechanisms, see Herbette and Ross (2023). See main text for references.

| System | Timing of PGE | How sexual karyotypes are established | Role of *doublesex* in sex determination |
| --- | --- | --- | --- |
| Dark-winged fungus gnats (Sciaridae) | Meiosis I | X elimination | No sex-specific splicing |
| Gall midges (Cecidomyiidae) | Meiosis I | X elimination | Not identified or characterised |
| Booklice (Psocodea) | Post-meiosis | Paternal genome silencing | Not identified or characterised |
| Parasitic lice (Phthiraptera) | Post-meiosis | Unknown | No sex-specific splicing |
| Scale insects (Neococcidae) | Meiosis II | Paternal genome silencing | No sex-specific splicing |
| Armoured scale insects (Diaspididae) | Early embryogenesis | Embryonic PGE | Not identified or characterised |
| Coffee berry borer beetle (*Hypothenemus*) | Meiosis I | Paternal genome silencing | Not identified or characterised |
| Globular springtails (Symphypleona) | Meiosis I | X elimination | Not identified or characterised |
| Predatory mites (Phytoseiidae) | Early embryogenesis | Embryonic PGE | Not identified or characterised |

In summary, our results highlight the potential for parent-offspring (and specifically maternal-zygotic) conflict over sex allocation in systems with asymmetric chromosome inheritance, with the magnitude of such conflicts varying depending on the demography and mating system. Under PGE, maternal and zygotic optima diverge, setting the stage for an evolutionary arms race that may play out at the molecular level in early development when sex is determined. While current evidence suggests that mothers usually dominate this process, convergent loss of canonical *doublesex* splicing in divergent taxa hints at a possible history of turnover. Resolving the extent of this conflict will require further functional and comparative work across a broader range of PGE systems. Such efforts may help to reveal not only how sex is determined in these unusual systems, but also how intragenomic conflict leaves lasting imprints on key developmental pathways.

## Acknowledgements and Funding

TJH is supported by the RIKEN Special Postdoctoral Researchers Program, and a RIKEN Incentive Grant. RBB is supported by a JSPS Postdoctoral Fellowship for research in Japan (PE) and the Howard Hughes Medical Institute (HHMI).

## Author contributions

TJH and RBB jointly developed the model, performed the analysis, and wrote the manuscript.

## Data and code availability statement

There is no associated data. Additional analysis can be found in the Supplementary Material.

