## Supplementary Material for "Parent-offspring conflict over sex determination in non-Mendelian systems"

#### 1 Life-cycle and genetics

##### 1.1 Genetics

We assume that all offspring initially have two gene copies, one of maternal-origin and one of paternal-origin. The following parameters control the transmission genetics. Offspring inherit their mother's maternal-origin copy with probability $\alpha$, and their father's paternal-origin copy with probability $\beta$. Standard Mendelian diploidy would correspond to $\alpha = 1/2$, $\beta = 1/2$, and paternal genome elimination would be $\alpha = 1/2$, $\beta = 0$. This is visualised in Figure 2a of the main text.

Individuals contain two gene copies which may influence the phenotype they express. We modulate the influence of these two genomes with the following parameters. For zygotes, the fraction of their phenotype controlled by their maternal-origin genes is $\rho_{\mathrm{Z}}$. For adult females (mothers) this is $\rho_{\mathrm{M}}$, and for adult males (fathers) this is $\rho_{\mathrm{M}}$. Thus in the case of equal expression $\rho_{\mathrm{i}} = 1/2$. If fathers silence their paternal-origin gene copy then $\rho_{\mathrm{F}} = 1$.

##### 1.2 Life cycle

We consider an infinite population size, divided into a series of patches; an infinite island model (Wright, 1931). Upon each patch, a large number of offspring are born, they then sexually differentiate into either males or females. Males then disperse from the patch. The probability an individual male remains on his natal patch is $h_{\mathrm{m}}$.

We then consider two life cycle variants, depending on whether females mate before, or after, they disperse. We will refer to these as Dispersal-Dispersal-Mating (DDM), and Dispersal-Mating-Dispersal (DMD) (Wild and

Taylor, 2004). The probability a females remains on her focal patch is $h_\mathrm{f}$. After both the mating and dispersal, mated females then compete for breeding spots upon each patch, produce a large number of offspring, and then die.

Within this life cycle a number of different mating systems are possible. Previously, both Pen (2006), and Wild & West (2009) used similar approaches to describe three types of mating systems: monogamy, polyandry, and polygyny. We use the following notation to describe these system. The probability that two offspring on a patch share the same mother is $A$. Given two offspring share the same mother, the probability that they share the same father is $B_i$. If offspring do not share the same mother, but their mothers mated on the same patch, then the probability that they share the same father is $B_p$.

With this notation monogamy corresponds to $B_i = 1$ and $B_p = 0$. Polyandry corresponds to $B_i = (1/M)$ and $B_p = 0$, where $M$ is the number of males that a female mates with. Polygyny corresponds to $B_i = 1$ and $B_p = 1$. Whilst we solve for arbitrary values of these parameters, we focus on these three cases for comparisons to previous work.

### 2 Consanguinities and relatedness

#### 2.1 Within and between juveniles

We compute the consanguinity between gene copies of maternal or paternal origin within and between juvenile individuals. Our general notation for relatedness is $r_{ij}^{\tau\kappa}$, where the lower indices $i,j \in \{m,p\}$ denote the parental origin of the two genes, the upper index $\tau \in \{Z,M,F\}$ identifies the controlling actor (zygote, mother, or father), and $\kappa \in \{S,P\}$ indicates whether the recipient is oneself (or one's own offspring) ($S$) or a patchmate juvenile ($P$). For example, the relatedness between the maternal-origin gene in a mother and the paternal-origin gene in a patch juvenile is $r_{mp}^{MP}$.

In the juvenile stage the controlling actor is the zygote ($\tau = Z$), and for notational simplicity we omit this superscript initially. Thus the four juvenile consanguinities we track are collected in the vector

$$\mathbf{R} = \begin{pmatrix} r_{mp}^{S} \\ r_{mm}^{P} \\ r_{mp}^{P} \\ r_{pp}^{P} \end{pmatrix}, \tag{S1}$$

where $r_{ij}^{S} = r_{ij}^{ZS}$ and $r_{ij}^{P} = r_{ij}^{ZP}$. Because individuals are diploid, we have $r_{ii}^{S} = 1$.

These consanguinities evolve according to the linear recursion

$$\mathbf{R}' = \mathbf{S}\mathbf{R} + \mathbf{C}, \tag{S2}$$

where $\mathbf{S}$ encodes the transmission and dispersal components of the life cycle, and $\mathbf{C}$ contains the corresponding probabilities of coalescence. We decompose these into genetic and demographic contributions,

$$\mathbf{S} = \mathbf{S}_g \circ \mathbf{S}_d, \qquad \mathbf{C} = \mathbf{C}_g \circ \mathbf{C}_d, \tag{S3}$$

where $\circ$ denotes elementwise multiplication.

The genetic transmission matrix is

$$\mathbf{S}_g = \begin{pmatrix} 0 & \alpha(1-\beta) & \alpha\beta + (1-\alpha)(1-\beta) & (1-\alpha)\beta \\ 2(1-\alpha)\alpha & \alpha^2 & 2(1-\alpha)\alpha & (1-\alpha)^2 \\ 0 & \alpha(1-\beta) & \alpha\beta + (1-\alpha)(1-\beta) & (1-\alpha)\beta \\ 2(1-\beta)\beta & (1-\beta)^2 & 2(1-\beta)\beta & \beta^2 \end{pmatrix}, \tag{S4}$$

and the corresponding coalescence vector is

$$\mathbf{C}_g = \begin{pmatrix} 0 \\ \alpha^2 + (1-\alpha)^2 \\ 0 \\ \beta^2 + (1-\beta)^2 \end{pmatrix}. \tag{S5}$$

The demographic matrices depend on the life cycle. For the DDM case,

$$\mathbf{S}_{DDM} = \begin{pmatrix} 0 & h_f h_m & h_f h_m & h_f h_m \\ A & (1-A)h_f^2 & (1-A)h_f^2 & (1-A)h_f^2 \\ 0 & h_f h_m & h_f h_m & h_f h_m \\ AB_i + (1-A)B_p & h_m^2\left(1 - AB_i - (1-A)B_p\right) & h_m^2\left(1 - AB_i - (1-A)B_p\right) & h_m^2\left(1 - AB_i - (1-A)B_p\right) \end{pmatrix}, \tag{S6}$$

with coalescence vector

$$\mathbf{C}_{DDM} = \begin{pmatrix} 0 \\ A \\ 0 \\ AB_i + (1-A)B_p \end{pmatrix}. \tag{S7}$$

For the DMD life cycle we obtain

$$\mathbf{S}_{DMD} = \begin{pmatrix} 0 & h_m & h_m & h_m \\ A & (1-A)h_f^2 & (1-A)h_f^2 & (1-A)h_f^2 \\ 0 & h_m Y & h_m Y & h_m Y \\ AB_i + (1-A)h_f^2 B_p & h_m^2 X & h_m^2 X & h_m^2 X \end{pmatrix}, \tag{S8}$$

where

$$\begin{aligned} Y &= A + (1-A)\,h_f^2, \\ X &= A(1-B_i) + (1-A)\,h_f^2(1-B_p). \end{aligned} \tag{S9}$$

The coalescence vector is

$$\mathbf{C}_{DMD} = \begin{pmatrix} 0 \\ A \\ 0 \\ AB_i + (1-A)h_f^2 B_p \end{pmatrix}. \tag{S10}$$

We solve these recursions for the equilibrium consanguinities, although we do not write the full expressions explicitly.

#### 2.2 Parents to offspring and patchmates

Next, we can use the above relatedness coefficients to compute the relatedness of parents to their offspring, and the other zygotes residing on the patch.

##### 2.2.1 Dispersal-Dispersal-Mating

From mothers to their own offspring:

$$\begin{aligned}
r_{mm}^{MS} &= \alpha + (1-\alpha)\, r_{mp}^{S},\\
r_{mp}^{MS} &= h_f h_m \Big[(1-\beta)\, r_{mm}^{P} + \beta\, r_{mp}^{P}\Big],\\
r_{pm}^{MS} &= \alpha\, r_{mp}^{S} + (1-\alpha),\\
r_{pp}^{MS} &= h_f h_m \Big[(1-\beta)\, r_{mp}^{P} + \beta\, r_{pp}^{P}\Big].
\end{aligned} \tag{S11}$$

From mothers to other zygotes on the patch:

$$\begin{aligned}
r_{mm}^{MP} &= A\Big[\alpha + (1-\alpha)\, r_{mp}^{S}\Big] + (1-A)\, h_f^2 \Big[\alpha\, r_{mm}^{P} + (1-\alpha)\, r_{mp}^{P}\Big],\\
r_{mp}^{MP} &= h_f h_m \Big[(1-\beta)\, r_{mm}^{P} + \beta\, r_{mp}^{P}\Big],\\
r_{pm}^{MP} &= A\Big[\alpha\, r_{mp}^{S} + (1-\alpha)\Big] + (1-A)\, h_f^2 \Big[\alpha\, r_{mp}^{P} + (1-\alpha)\, r_{pp}^{P}\Big],\\
r_{pp}^{MP} &= h_f h_m \Big[(1-\beta)\, r_{mp}^{P} + \beta\, r_{pp}^{P}\Big].
\end{aligned} \tag{S12}$$

From fathers to their own offspring:

$$\begin{aligned}
r_{mm}^{FS} &= h_f h_m \Big[\alpha\, r_{mm}^{P} + (1-\alpha)\, r_{mp}^{P}\Big],\\
r_{mp}^{FS} &= (1-\beta) + \beta\, r_{mp}^{S},\\
r_{pm}^{FS} &= h_f h_m \Big[\alpha\, r_{mp}^{P} + (1-\alpha)\, r_{pp}^{P}\Big],\\
r_{pp}^{FS} &= (1-\beta)\, r_{mp}^{S} + \beta.
\end{aligned} \tag{S13}$$

From fathers to the other zygotes on their offspring's patch:

$$\begin{aligned}
r_{mm}^{FP} &= h_f h_m \Big[\alpha\, r_{mm}^{P} + (1-\alpha)\, r_{mp}^{P}\Big],\\
r_{mp}^{FP} &= \big[AB_i + (1-A)B_p\big]\big[(1-\beta) + \beta\, r_{mp}^{S}\big]\\
&\quad + \Big(1 - \big[AB_i + (1-A)B_p\big]\Big) h_m^2 \Big[(1-\beta)\, r_{mm}^{P} + \beta\, r_{mp}^{P}\Big],\\
r_{pm}^{FP} &= h_f h_m \Big[\alpha\, r_{mp}^{P} + (1-\alpha)\, r_{pp}^{P}\Big],\\
r_{pp}^{FP} &= \big[AB_i + (1-A)B_p\big]\big[(1-\beta)\, r_{mp}^{S} + \beta\big]\\
&\quad + \Big(1 - \big[AB_i + (1-A)B_p\big]\Big) h_m^2 \Big[(1-\beta)\, r_{mp}^{P} + \beta\, r_{pp}^{P}\Big].
\end{aligned} \tag{S14}$$

###### 2.2.2 Dispersal-Mating-Dispersal

Mothers to their own offspring:

$$\begin{aligned}
r_{mm}^{MS} &= \alpha + (1-\alpha)\, r_{mp}^{S},\\
r_{mp}^{MS} &= h_m \Big[(1-\beta)\, r_{mm}^{P} + \beta\, r_{mp}^{P}\Big],\\
r_{pm}^{MS} &= \alpha\, r_{mp}^{S} + (1-\alpha),\\
r_{pp}^{MS} &= h_m \Big[(1-\beta)\, r_{mp}^{P} + \beta\, r_{pp}^{P}\Big].
\end{aligned} \tag{S15}$$

Mothers to the other zygotes on their offspring's patch:

$$\begin{aligned}
r_{mm}^{MP} &= A\Big[\alpha + (1-\alpha)\, r_{mp}^{S}\Big] + (1-A)\, h_f^2 \Big[\alpha\, r_{mm}^{P} + (1-\alpha)\, r_{mp}^{P}\Big],\\
r_{mp}^{MP} &= \big(Ah_m + (1-A) h_f^2 h_m\big)\big[(1-\beta)\, r_{mm}^{P} + \beta\, r_{mp}^{P}\big],\\
r_{pm}^{MP} &= A\Big[\alpha\, r_{mp}^{S} + (1-\alpha)\Big] + (1-A)\, h_f^2 \Big[\alpha\, r_{mp}^{P} + (1-\alpha)\, r_{pp}^{P}\Big],\\
r_{pp}^{MP} &= \big(Ah_m + (1-A) h_f^2 h_m\big)\big[(1-\beta)\, r_{mp}^{P} + \beta\, r_{pp}^{P}\big].
\end{aligned} \tag{S16}$$

Fathers to their own offspring:

$$\begin{aligned}
r_{mm}^{FS} &= h_m \Big[\alpha\, r_{mm}^{P} + (1-\alpha)\, r_{mp}^{P}\Big],\\
r_{mp}^{FS} &= (1-\beta) + \beta\, r_{mp}^{S},\\
r_{pm}^{FS} &= h_m \Big[\alpha\, r_{mp}^{P} + (1-\alpha)\, r_{pp}^{P}\Big],\\
r_{pp}^{FS} &= (1-\beta)\, r_{mp}^{S} + \beta.
\end{aligned} \tag{S17}$$

Fathers to the other zygotes on their offspring's patch:

$$
\begin{aligned}
r^{FP}_{mm} &= \big(h_m(A+(1-A)h_f^2)\big)\big[\alpha\, r^P_{mm} + (1-\alpha)\, r^P_{mp}\big],\\
r^{FP}_{mp} &= \big(AB_i + (1-A)B_p h_f^2\big)\big[(1-\beta) + \beta\, r^S_{mp}\big]\\
&\quad + \Big(A(1-B_i)h_m^2 + (1-A)(1-B_p)h_f^2 h_m^2\Big)\big[(1-\beta)\, r^P_{mm} + \beta\, r^P_{mp}\big],\\
r^{FP}_{pm} &= \big(h_m(A+(1-A)h_f^2)\big)\big[\alpha\, r^P_{mp} + (1-\alpha)\, r^P_{pp}\big],\\
r^{FP}_{pp} &= \big(AB_i + (1-A)B_p h_f^2\big)\big[(1-\beta)\, r^S_{mp} + \beta\big]\\
&\quad + \Big(A(1-B_i)h_m^2 + (1-A)(1-B_p)h_f^2 h_m^2\Big)\big[(1-\beta)\, r^P_{mp} + \beta\, r^P_{pp}\big].
\end{aligned}
\tag{S18}
$$

### 3 Reproductive values

We can compute the class reproductive values, i.e. the expected long term contribution of one class of individuals today to the ancestry of the population. If we census as juveniles, then we have two set of genes of interest, maternal-origin genes and paternal-origin genes. We write out the following transition matrix, which describes the probability that a gene found in state $i$ in this generation was in state $j$ in the previous one. With the transmission genetics described earlier:

$$
\begin{pmatrix} c_{\mathbb{m}} & c_{\mathbb{p}} \end{pmatrix} = \begin{pmatrix} c_{\mathbb{m}} & c_{\mathbb{p}} \end{pmatrix} \begin{pmatrix} \alpha & 1-\alpha \\ 1-\beta & \beta \end{pmatrix} = \begin{pmatrix} \frac{1-\beta}{2-\alpha-\beta} & \frac{1-\alpha}{2-\alpha-\beta} \end{pmatrix} \tag{S19}
$$

From the maternal-origin and paternal-origin genes, we can then write out the class reproductive values of these gene sets in males and females. As the value of males are equivalent to the paternal-origin genes (as all of these genes derive from males) $c_{\mathbb{p}} = c_m$. Then the relative value of the maternal-origin and paternal-origin genes are given by their share in this, and so:

$$
c_{f_{\mathbb{m}}} = \alpha c_{\mathbb{m}} = \frac{\alpha(1-\beta)}{2-\alpha-\beta} \tag{S20}
$$

$$
c_{f_{\mathbb{p}}} = (1-\alpha) c_{\mathbb{m}} = \frac{(1-\alpha)(1-\beta)}{2-\alpha-\beta} \tag{S21}
$$

$$
c_{m_{\mathbb{m}}} = (1-\beta) c_{\mathbb{p}} = \frac{(1-\beta)(1-\alpha)}{2-\alpha-\beta} \tag{S22}
$$

$$
c_{m_{\mathbb{p}}} = \beta c_{\mathbb{p}} = \frac{\beta(1-\alpha)}{2-\alpha-\beta} \tag{S23}
$$

### 4 Marginal inclusive fitness changes

We can now write the change in inclusive fitness as a function of relatedness and class reproductive values. We consider an offspring controlling the sex ratio decision, and a small deviation $x = z + \delta$ in its sex allocation strategy, with $\delta$ small. We ask whether this mutant strategy can invade from rarity.

A change in sex allocation is favoured if, on average, it increases the inclusive fitness of the actor. Under offspring control there are two types of actor: maternal-origin genes in offspring and paternal-origin genes in offspring, present at relative frequencies $u_{\mathbb{m}} = 1/2$ and $u_{\mathbb{p}} = 1/2$.

For a maternal-origin actor, the change in inclusive fitness from an increased allocation to males is

$$\begin{aligned}\Delta\Omega_{\mathbb{m}} = \rho\Bigg( &+ r^{ZS}_{mm}\left(\frac{c_{m_{\mathbb{m}}}}{u_{m_{\mathbb{m}}}} - \frac{c_{f_{\mathbb{m}}}}{u_{f_{\mathbb{m}}}}\right) + r^{ZS}_{mp}\left(\frac{c_{m_{\mathbb{p}}}}{u_{m_{\mathbb{p}}}} - \frac{c_{f_{\mathbb{p}}}}{u_{f_{\mathbb{p}}}}\right) \\ &- r^{ZP}_{mm}\left(a_m \frac{c_{m_{\mathbb{m}}}}{u_{m_{\mathbb{m}}}} - a_f \frac{c_{f_{\mathbb{m}}}}{u_{f_{\mathbb{m}}}} + a_{\text{fm}} \frac{u_{m_{\mathbb{m}}}}{u_{f_{\mathbb{m}}}} \frac{c_{m_{\mathbb{m}}}}{u_{m_{\mathbb{m}}}}\right) \\ &- r^{ZP}_{mp}\left(a_m \frac{c_{m_{\mathbb{p}}}}{u_{m_{\mathbb{p}}}} - a_f \frac{c_{f_{\mathbb{p}}}}{u_{f_{\mathbb{p}}}} + a_{\text{fm}} \frac{u_{m_{\mathbb{p}}}}{u_{f_{\mathbb{p}}}} \frac{c_{m_{\mathbb{p}}}}{u_{m_{\mathbb{p}}}}\right)\Bigg)\end{aligned} \tag{S24}$$

The change in inclusive fitness for a paternal-origin gene is

$$\begin{aligned}\Delta\Omega_{\mathbb{p}} = (1-\rho)\Bigg( &+ r^{ZS}_{pm}\left(\frac{c_{m_{\mathbb{m}}}}{u_{m_{\mathbb{m}}}} - \frac{c_{f_{\mathbb{m}}}}{u_{f_{\mathbb{m}}}}\right) + r^{ZS}_{pp}\left(\frac{c_{m_{\mathbb{p}}}}{u_{m_{\mathbb{p}}}} - \frac{c_{f_{\mathbb{p}}}}{u_{f_{\mathbb{p}}}}\right) \\ &- r^{ZP}_{pm}\left(a_m \frac{c_{m_{\mathbb{m}}}}{u_{m_{\mathbb{m}}}} - a_f \frac{c_{f_{\mathbb{m}}}}{u_{f_{\mathbb{m}}}} + a_{\text{fm}} \frac{u_{m_{\mathbb{m}}}}{u_{f_{\mathbb{m}}}} \frac{c_{m_{\mathbb{m}}}}{u_{m_{\mathbb{m}}}}\right) \\ &- r^{ZP}_{pp}\left(a_m \frac{c_{m_{\mathbb{p}}}}{u_{m_{\mathbb{p}}}} - a_f \frac{c_{f_{\mathbb{p}}}}{u_{f_{\mathbb{p}}}} + a_{\text{fm}} \frac{u_{m_{\mathbb{p}}}}{u_{f_{\mathbb{p}}}} \frac{c_{m_{\mathbb{p}}}}{u_{m_{\mathbb{p}}}}\right)\Bigg)\end{aligned} \tag{S25}$$

Inspecting these expressions, we can identify three main components of the inclusive-fitness effect of a marginal shift in sex allocation.

(1) A direct change in the numbers of sons and daughters produced: an increase of $\delta$ males and a decrease of $\delta$ females. These individuals contribute $c_m/u_m$ and $c_f/u_f$, respectively, to the actor's inclusive fitness (with the appropriate parent-of-origin subscripts in the class indices).

(2) A change in the intensity of sex-specific local competition among same-sex patchmates. Increasing the production of males intensifies competition among males and relaxes competition among females, with marginal effects $-a_m \frac{c_m}{u_m}$ and$-a_f \frac{c_f}{u_f}$, again applied to the relevant maternal- and paternal-origin classes.

(3) A cross-sex effect on the number of mating opportunities for males. Changing the local sex ratio alters the availability of mates, with a marginal effect $-a_{\text{fm}} \frac{u_m}{u_f} \frac{c_m}{u_m}$, reflecting the effect of the changing number of mates upon the reproductive value of local males.

The values of the different scales of competition under the two demographic regimes considered - DDM and DMD - can be seen inS1.

| | $a_f$ | $a_m$ | $a_{fm}$ |
|---|---|---|---|
| DDM | $h_f^2$ | $h_m^2$ | 0 |
| DMD | $h_f^2$ | $h_m^2$ | $(1-h_f^2)\,h_m$ |

Table S1: Scales of competition ($a$'s) under the two demographic regimes analysed (DDM and DMD).

We aggregate the total change in inclusive fitness over the two types of actor: $\Delta\Omega = u_{\mathbb{m}}\Delta\Omega_{\mathbb{m}} + u_{\mathbb{p}}\Delta\Omega_{\mathbb{p}}$, Multiplying the inclusive fitness change of these two actors by their relative frequency in the population. In this case $u_{\mathbb{m}} = u_{\mathbb{p}} = 1/2$. A mutant allocation strategy $x = z + \delta$ can invade when

$$\Delta\Omega > 0. \tag{S26}$$

The candidate singular strategy $z^*$ is obtained by solving $\Delta\Omega = 0$ for $z$, and we determine whether $z^*$ is an evolutionary stable sex allocation (fitness maximum) or a fitness minimum by examining the sign of the marginal fitness change around $z^*$ (Taylor, 1996).

The inclusive–fitness effect under offspring control, $\Delta\Omega_Z$, provides a template for all actor classes. The corresponding expressions for maternal- and paternal-control scenarios are obtained by substituting the appropriate actor-specific relatedness coefficients $r_{ij}^{\tau\kappa}$ and phenotypic control parameters $\rho_\tau$ (with $\tau \in \{M, F\}$) into the above equations. Results for the different demographic scenarios are presented below in both tables and graphically.

### 5 Tables and Figures

| **Genetic system** | **Mating structure** | **Offspring** | **Mothers** | **Fathers** |
|---|---|---|---|---|
| Diploidy | Monogamy | $\frac{3N+1}{6N}$ | $\frac{N(3N+1)}{6N^2-3N+1}$ | $\frac{N}{2N-1}$ |
| | Polyandry | $\frac{M(3N+1)}{6MN+M-1}$ | $\frac{MN(3N+1)}{6MN^2-(2M+1)N+1}$ | $\frac{MN}{2MN-1}$ |
| | Polygamy | $\frac{3N+1}{5N+1}$ | $\frac{3N+1}{5N-1}$ | 1 |
| PGE | Monogamy | $\frac{3N+1}{9N-3}$ | $\frac{N(3N+1)}{6N^2-4N+2}$ | 0 |
| | Polyandry | $\frac{M(3N+1)}{9MN+M-4}$ | $\frac{MN(3N+1)}{6MN^2-2(M+1)N+2}$ | 0 |
| | Polygamy | $\frac{3N+1}{5N+1}$ | $\frac{3+\frac{1}{N}}{4}$ | No optimum |

Table S2: Optimal sex ratios under different genetic systems and mating structures in a local resource competition model, where males always disperse ($h_m = 0$) and females never disperse ($h_f = 1$).This is identical under the DDM and DMD life cycles. $N$ is the number of females on the patch, and $M$ is the number of males each female mates with. All results presented are for joint control ($\rho = 1/2$)

| Genetic system | Mating structure | Offspring | Mothers | Fathers |
|---|---|---|---|---|
| Diploidy | Monogamy | $\frac{1}{2}-\frac{1}{6N}$ | $\frac{N-1}{2N-1}$ | $1-\frac{N(3N+1)}{6N^2-3N+1}$ |
| | Polyandry | $\frac{M(3N-1)}{1+M(6N-1)}$ | $\frac{N-1}{2N-1}$ | $\frac{(3N-1)(MN-1)}{6MN^2-(2+M)N+1}$ |
| | Polygyny | $\frac{3N-1}{7N-1}$ | $\frac{N-1}{2N-1}$ | 0 |
| PGE | Monogamy | $\frac{2N-1}{6N-1}$ | $\frac{N-1}{2N-1}$ | 0 |
| | Polyandry | $\frac{2N-1}{6N-1}$ | $\frac{N-1}{2N-1}$ | 0 |
| | Polygyny | $\frac{2N-1}{6N-1}$ | $\frac{N-1}{2N-1}$ | 0 |

Table S3: Optimal sex ratios under different genetic systems and mating structures in a local mate competition model, where males do not disperse ($h_m = 1$) and females always disperse before mating ($h_f = 0$), i.e. the DDM life cycle. $N$ is the number of females on the patch, and $M$ is the number of males each female mates with. All results presented are for joint control ($\rho = 1/2$). Note that under PGE, the mating system exerts no impact upon results.

| Genetic system | Mating structure | Offspring | Mothers | Fathers |
|---|---|---|---|---|
| Diploidy | Monogamy | $\frac{N-1}{2N-1}$ | $\frac{N-1}{2N}$ | $\frac{N-1}{2N}$ |
| | Polyandry (large $M$) | $\frac{(2N-1)^2}{8N^2-6N+2}$ | $\frac{N-1}{2N}$ | $\frac{1}{2}-\frac{1}{4N^2-2N+1}$ |
| | Polygyny | $\frac{N-1}{2N-1}$ | $\frac{N-1}{2N}$ | $\frac{N-1}{2N}$ |
| PGE | Monogamy | $\frac{2(N-1)}{6N-3}$ | $\frac{(N-1)(2N-1)}{N(4N-1)}$ | $\frac{(N-1)(3N-1)}{2N(2N^2+2N-1)}$ |
| | Polyandry (large $M$) | $\frac{2(N-1)}{6N-3}$ | $\frac{(N-1)(2N-1)}{N(4N-1)}$ | $\frac{2N^3+N^2-4N+1}{2N(4N^3+3N-1)}$ |
| | Polygyny | $\frac{2(N-1)}{6N-3}$ | $\frac{(N-1)(2N-1)}{N(4N-1)}$ | $\frac{(N-1)(3N-1)}{2N(2N^2+2N-1)}$ |

Table S4: Optimal sex ratios under different genetic systems and mating structures in a local mate competition model, where males do not disperse ($h_m = 1$) and females always disperse after mating ($h_f = 0$), i.e. the DMD life cycle. $N$ is the number of females on the patch. Results for arbitrary $M$ are unwieldy, and therefore we present a large $M$ approximation here ($M \to \infty$). All results presented are for joint control ($\rho = 1/2$).

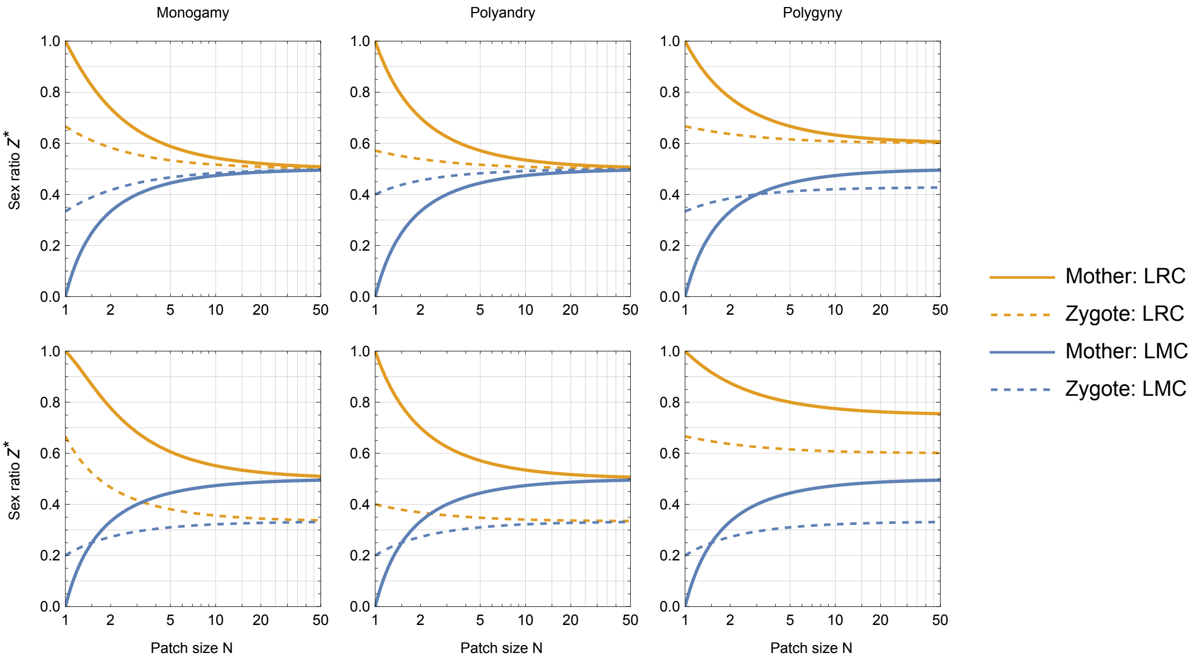


Figure S1: **Optimal sex ratio under a Dispersal–Dispersal–Mating life cycle.** Analytical solutions for optimal sex ratios under different genetic systems—diploidy (top row) and paternal genome elimination (bottom row)—across mating systems(monogamy, polyandry, and polygyny) and controlling parties (mothers versus offspring).Polyandry cases were derived under the large-$M$ approximation.

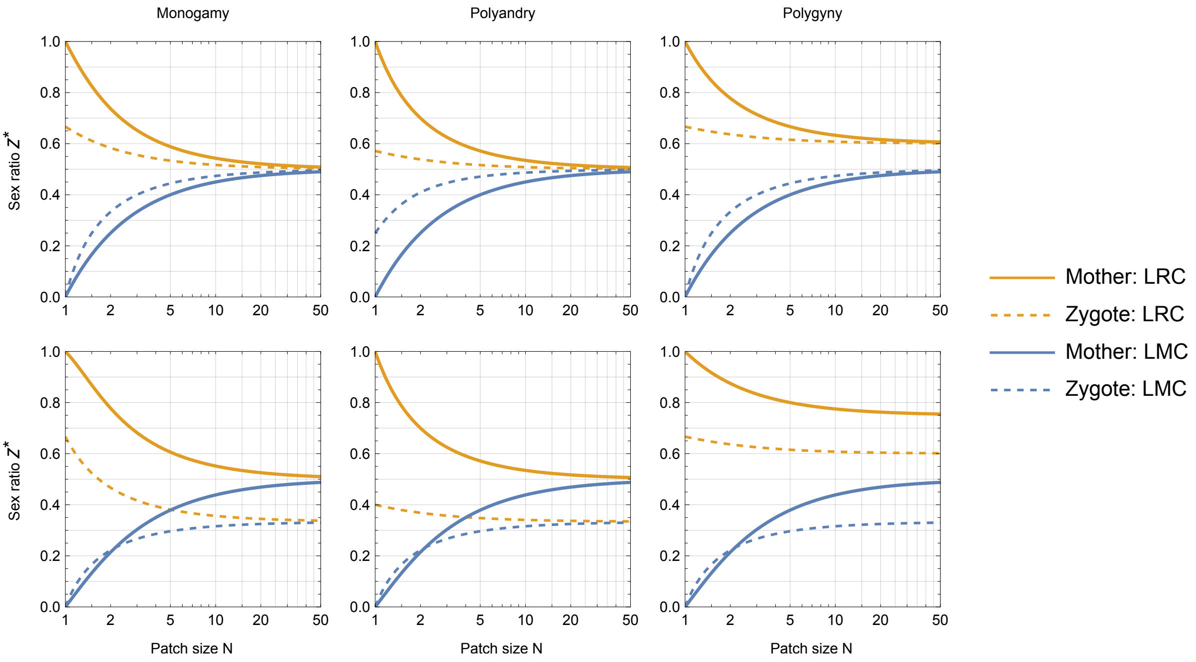


Figure S2: **Optimal sex ratio under a Dispersal–Mating–Dispersal life cycle.** Analytical solutions for optimal sex ratios under different genetic systems—diploidy (top row) and paternal genome elimination (bottom row)—across mating systems(monogamy, polyandry, and polygyny) and controlling parties (mothers versus offspring).Polyandry cases were derived under the large-$M$ approximation.
